# PlantRing: A high-throughput wearable sensor system for decoding plant growth, water relations and innovating irrigation

**DOI:** 10.1101/2024.11.29.625988

**Authors:** Ting Sun, Chenze Lu, Zheng Shi, Mei Zou, Peng Bi, Xiaodong Xu, Qiguang Xie, Rujia Jiang, Yunxiu Liu, Rui Cheng, Wenzhao Xu, Yingying Zhang, Pei Xu

## Abstract

The combination of flexible electronics and plant science has generated various plant-wearable sensors, yet challenges persist in their applications in real-world agriculture, particularly in high-throughput settings. Overcoming the trade-off between sensing sensitivity and range, adapting them to a wide range of crop types, and bridging the gap between sensor measurements and biological understandings remain the primary obstacles. Here we introduce PlantRing, an innovative, nano-flexible sensing system designed to address the aforementioned challenges. PlantRing employs bio-sourced carbonized silk georgette as the strain sensing material, offering exceptional resolution (tensile deformation: < 100 μm), stretchability (tensile strain up to 100 %), and remarkable durability (season long), exceeding existing plant strain sensors. PlantRing effectively monitors plant growth and water status, by measuring organ circumference dynamics, performing reliably under harsh conditions and being adaptable to a wide range of plants. Applying PlantRing to study fruit cracking in tomato and watermelon reveals novel hydraulic mechanism, characterized by genotype-specific excess sap flow within the plant to fruiting branches. Its high-throughput application enabled large-scale quantification of stomatal sensitivity to soil drought, a traditionally difficult-to-phenotype trait, facilitating drought tolerant germplasm selection. Combing PlantRing with soybean mutant led to the discovery of a potential novel function of the *GmLNK2* circadian clock gene in stomatal regulation. More practically, integrating PlantRing into feedback irrigation achieves simultaneous water conservation and quality improvement, signifying a paradigm shift from experience- or environment-based to plant-based feedback control. Collectively, PlantRing represents a groundbreaking tool ready to revolutionize botanical studies, agriculture, and forestry.

## Introduction

The ability to trace crop status accurately, dynamically, and continuously amidst environmental fluctuations provides a foundation for adequate interventions to improve yield and quality [1]. While various technologies have proven their feasibility in plant growth and health monitoring, practical agricultural implementation demands a seamless extension of such capabilities from individual to high-throughput levels and from controlled environments to natural settings [2]. Currently, commercially available high-throughput crop monitoring systems rely on optical systems (RGB, multispectral, and hyperspectral) or lysimetric arrays [3]. However, the former exhibits low spatial resolution and a limited capacity to monitor functional physiological traits compared to structural traits [4]. The latter is restricted by its immovability, necessity for specific cultivation containers, and high cost [5, 6].

The recent integration of nanotechnology and electronic interfaces has generated plant-wearable sensors, which by function can be categorized as sensors for the microclimate on plant surfaces, volatile organic compounds, organ growth, and intrinsic sap flow [7, 8]. They are crafted from diverse nanomaterials, such as chitin-based water ink [9], laser-induced graphene [10], carbon nanotube/graphite [11], and polyimide [12], which can be worn or printed directly on leaves, stems, or fruits, enabling non-invasive, real-time monitoring [7, 13]. However, several significant challenges have hindered the entry of plant wearables into the market and their use in agriculture. One primary challenge is the trade-off between sensing sensitivity and range, which is particularly problematic in thermal anisotropy-based sensors [14]. Additionally, stability and robustness of the wearable sensors must be confirmed in harsh environments over long growth seasons, along with readiness for high-throughput deployment on a wide range of crop types [15, 16]. As such, the successful transition of wearable sensors from laboratory demonstrations to practical applications in agriculture remains an ongoing journey. Furthermore, a significant gap exists in translating the sensor data into a physiological understanding and practical insights for agriculture. For example, the diurnal plant stem diameter variation (SDV), caused by radial shrinkage due to plant transpiration and radial expansion due to water uptake and growth [17], has long been considered a sensitive indicator of plant health under normal and stressed conditions [18, 19]. It is also a promising parameter for feedback irrigation [20]. Unfortunately, despite advancements in sensor technology enabling more effective SDV monitoring, a comprehensive understanding of the linkage between plant function and SDV remains nascent, and this gap in understanding has led to the integration of these sophisticated sensors into intelligent irrigation systems lagging behind.

Among the various types of wearable sensors, strain sensors convert mechanical deformations into changes in electrical characteristics such as resistance or capacitance and constitute a mainstream category of current plant wearables [21]. Employing biomaterials, we have developed a series of strain sensors with excellent biocompatibility and capacity [22, 23]. Our previous study established the technical excellence of a carbonized silk georgette-based strain sensor, demonstrating large stretchability (>100% strain), ultrahigh sensitivity (average gauge factor of 29.7 within a 40% strain and 173.0 for a strain of 60–100%), ultralow detection limit (0.01% strain), high durability and stability (10,000 stretching cycles at 100% strain), and fast response (<70 ms) [24]. Motivated by the need for robust and commercially viable plant wearables for agricultural use, here we present the high-throughput PlantRing system. It simultaneously achieves an low detection limit (0.1 mm) and a wide sensing stain range of up to 100% strain, overperforming existing strain sensors [25-27]. It maintains stability and robustness against common interferences in agriculture, including temperature, wind, and rain, without compromising plant viability over season-long applications. With the aid of PlantRing, we gained novel biological insights into the mechanism of fruit cracking and achieved quantification of the stomatal sensitivity to soil drought, a key trait for drought adaptation that has traditionally been challenging to phenotype, in a high-throughput manner. Additionally, we realized simultaneous water savings and quality enhancement of tomato fruits through the implementation of PlantRing-based feedback irrigation, positioning it as a potential game-changer in smart agriculture. To the best of our knowledge, PlantRing is currently the only economically viable plant wearable sensor system with proven versatile use in high-throughput, real-world agricultural applications.

## Results

### Design and fabrication of the PlantRing system

The PlantRing system comprised a sensor unit, waterproof wireless communication unit, and cloud-based software terminal designed for remote control, real-time data acquisition, display, and analysis (Fig. 1). Each sensor unit had a silk-georgette-based strain sensor that transformed mechanical deformations into resistance changes. This strain sensor featured a flexible and stretchable elastomer film-encapsulated carbonized commercial silk georgette comprising yarns made of natural silkworm silk fibers (tens of microns in diameter), distinguishing it from other documented plant sensors (Fig. 1a, S1 Table). The sensor length is customizable according to specific demands, rendering it versatile. Three sensor types, weighing only 6.7-21.7 g, have been manufactured based on their original length: 6 cm for measurement of stems (e.g. tomato and beans), and 12 and 30 cm for fruits of different sizes. The strain sensor was connected to a data logger equipped with two U-shaped handles for convenient attachment to the plant stems or support structures using automated cable ties. It included a printed circuit board containing a microprocessor chip and integrated with a commercial temperature and humidity sensor for environmental monitoring and data compensation. The plastic shells at both ends of the sensor serve as a flexible clip, allowing adjustment of the sensor length that wraps around the plant organ under examination (Fig. 1b, S1 Fig.). The microprocessor, equipped with an Analog-to-Digital Converter (ADC) module, processed physical deformation signals from the strain sensor through an operational amplifier circuit module (Fig. 1d, S2 Fig.). The data logger established communication with the gateway by using 2.4G RF technology, and the gateway used 4G/5G networks to transmit data or issue operational commands. The data logger has a minimum data transmission interval of 1 second and its transmission range covers a straight-line distance of up to 60 m. The sensing data were transmitted to a cloud sever hosted by Alibaba for remote monitoring, management and storage, accessible by both computer and smartphone (S3 Fig.). The system had low power consumption, allowing its continuous operation for at least three months with a rechargeable battery. Owing to the streamlined manipulation procedure, cost-effectiveness, light weight, and durability, our system is excellently suited for large-scale deployment, both in lab and greenhouse or field cultivation conditions.

**Figure 1.**
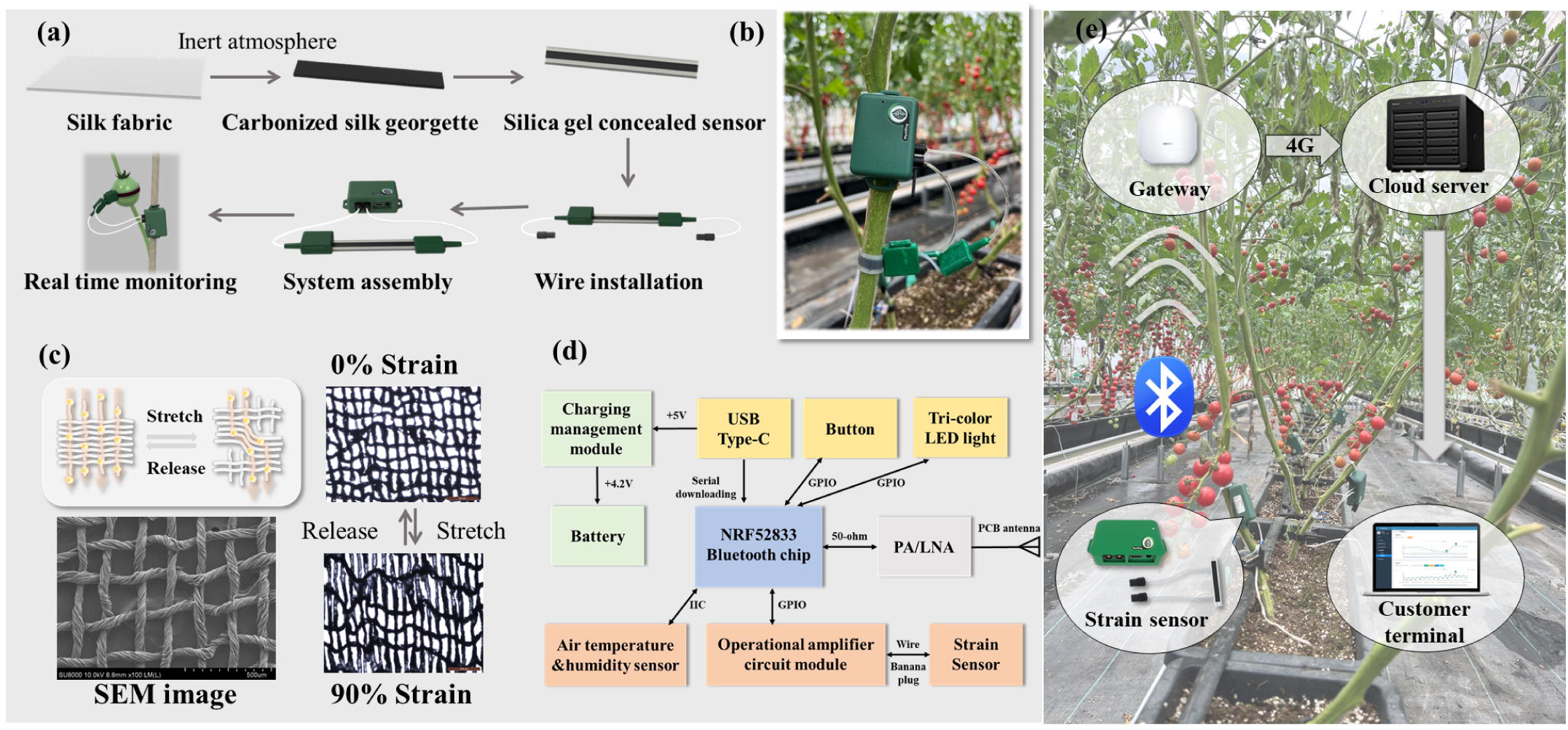
Design and fabrication of the PlantRing system. (a) The fabrication and assembly procedure of PlantRing. Silk fabric is carbonized under an inert atmosphere and sealed with silica gel. After installing the wire and data logger, the system is ready for real-time monitoring. (b) Image illustrating PlantRing installed on the stem of a tomato plant. (c) The principle of signal generation. The carbonized fabric displays different micromorphology when stretched, resulting in a change in resistant value. (d) The circuit design of PCB board in the data logger. The board is integrated with the NRF52833 Bluetooth chip to allow wireless transmission. (e) Demonstration of the overall concept of PlantRing system in agricultural use. The signal is wirelessly transmitted to a cloud server through a gateway, which provides real-time data to the user end for analysis or decision making.

### PlantRing system characterization

We tested the sensor performance in agricultural settings. Each sensor unit was calibrated before use by stretching it consistently to create a strain-to-AD signal response curve (S4 Fig.), which fit a linear function. A strong linear correlation (R^2^> 99.7%) for deformation was observed when the sensor was elongated within the range of 0-100% strain (Fig. 2a). Despite the difference in length, three sensor types all displayed excellent resolution (< 0.1 mm), repeatability (coefficient of variation < 0.68 %) and accuracy (relative error within the range of ±0.65%) (S2 Table).

**Figure 2.**
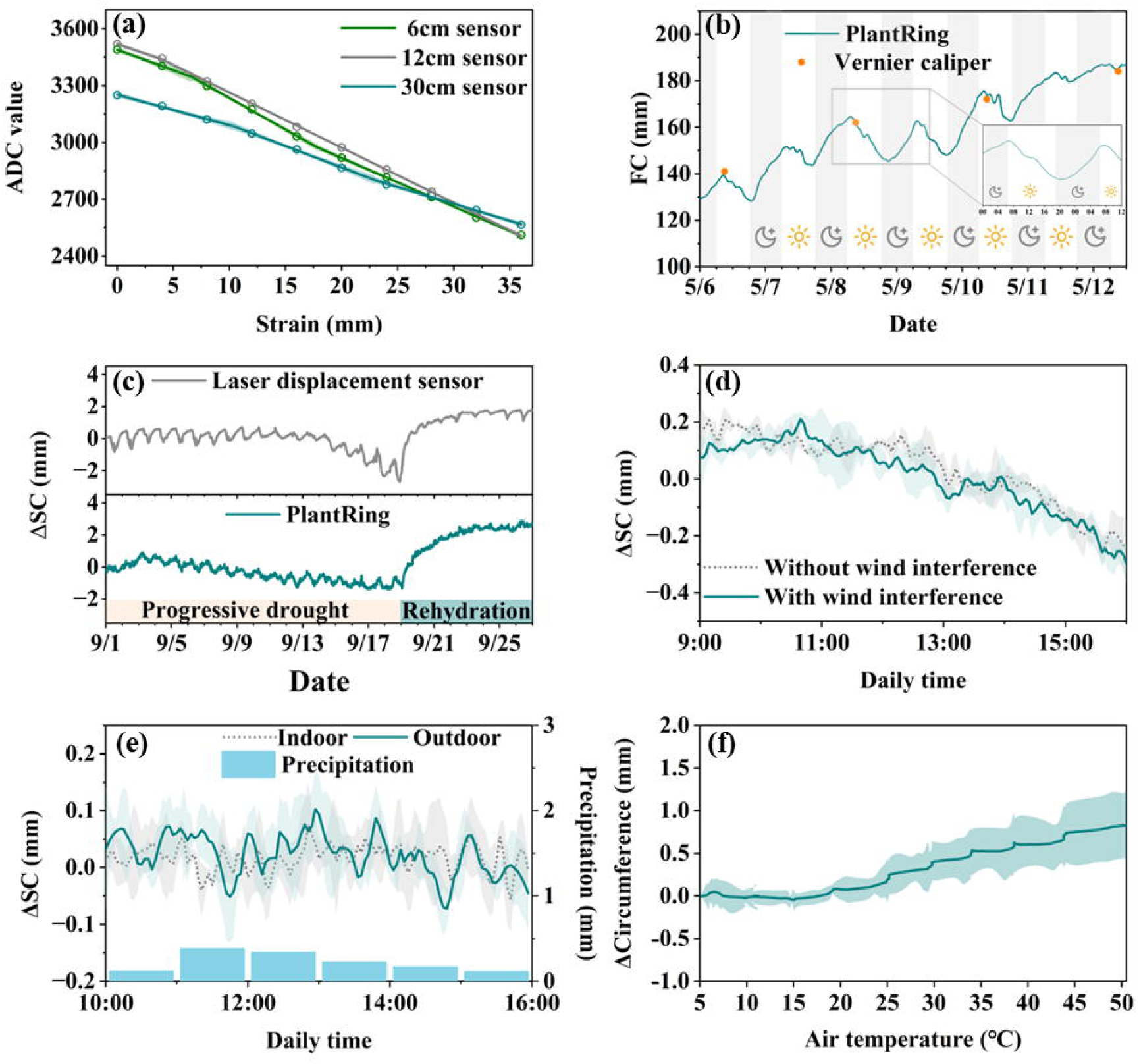
Characterization of the PlantRing system. Data are represented as mean ±SD. (a) Linear fit of the signal obtained from three types of sensor at different levels of strain. All sensors displayed good linearity in signal (R^2^>99.7%). The slope is recorded and written into the PCB of data logger for signal correction, which calibrates the patch difference of sensors during fabrication (b) Real-time fruit circumference (FC) measured with PlantRing in comparison with results obtained from a Vernier caliper. (c) The variation in stem circumference (ΔSC) obtained from diameters measured by laser displacement sensors (gray line) compared to ΔSC measured by PlantRing (blue line), illustrating the response to gradual soil drought and subsequent rehydration. The ΔSC value reset to zero at 0:00 on September 1^st^, 2023. (d) The ΔSC recorded by PlantRing for two groups of tomato plants: one with (blue line and shading) and one without (gray dotted line and shading) artificial airflow. The line represents the mean of three biological replicates, while the shaded area shows the variation between replicates. The ΔSC value reset to zero at 0:00 on November 28^th^, 2023. (e) The ΔSC recorded by PlantRing for two groups of tomato plants: one grown outdoors (blue line and shading) experiencing natural rain and wind interferences simultaneously, and one cultivated indoors in a greenhouse (gray dotted line and shading). The line represents the mean of four biological replicates, while the shaded area shows the variation between replicates. The ΔSC value reset to zero at 0:00 on December 4^th^, 2023. Hourly precipitation is shown as a bar graph at the bottom of the figure. (f) Circumference variation recorded by PlantRing when placed on a quartz glass rod, responding to temperatures of 5 and 50 °C. The gray and green lines depict, respectively, the average circumference variation without temperature calibration and after temperature calibration. The gray/green shading indicates the range of variation across three different sensors. Calibration was conducted utilizing temperature response data from 10 sensors, employing a quadratic function for temperature adjustment.

We then examined the functionality of the PlantRing system for measuring organ circumference. This entailed placing the strain sensors on the fruits and at the base of the stems of greenhouse-cultivated tomato plants. During the rapid growth phase of fruits, PlantRing continuously and accurately measured fruit circumference (FC) expansions, demonstrating a high level of agreement with the circumference values derived from diameters manually measured using calipers every three days (RMSE=1.2 mm, relative to fruit FC of 140-180 mm, Fig. 2b). The PlantRing data also revealed a trend of FC expansion over time due to fruit growth, along with a clear daily pattern of FC increasing during the day and decreasing at night (Fig. 2b), aligning with the measurement of nocturnal vine sap flow in watermelon through thermal-based sensors [14]. Likewise, tomato plants subjected to progressive soil drought and rehydration showed a diurnal fluctuation in stem diameter and circumference (SC), reflecting diurnal redistribution of stem water reserves influenced by daytime transpiration (Tr) and nighttime root water uptake (Fig. 2c) [28]. Nevertheless, a trend of SC reduction over extended droughts was recorded and was promptly reversed upon re-watering (Fig. 2c). This complex yet subtle phenomenon was validated by parallel measurements on the same plants using high-cost commercial laser displacement sensors (Pearson correlation coefficient = 0.87), highlighting the excellence of the PlantRing as a more economic and robust phenotyping tool (Fig. 2c, S5 Fig.).

Physical movements and microenvironmental changes (e.g. leaf surface temperature) induced by wind or rain pose challenges that limit the application of many existing wearable sensors under realistic agricultural conditions. As illustrated in Fig. 2d, the PlantRing-measured SC variation (ΔSC) of tomato plants exposed to artificial airflow (3.5 m/s) showed no significant difference from that of the control plants (*P*>0.05) (S1 Movie). In the presence of concurrent natural rain (Daily precipitation = 2.5 mm) and wind interferences (Daily mean wind speed = 1.8 m/s), PlantRing also operated normally (S2 Movie). In Fig. 2e, no significant difference (*P*>0.05) was observed in ΔSC between the indoor and outdoor groups, along with only slight change in ΔSC due to low transpirations under the low solar radiation of the day (150-251 μmol m^-2^ s^-1^ between 10:00 and 16:00). To examine the effect of temperature, we used a stimulation experiment employing a sensor mounted on a quartz glass rod, which had a very low thermal expansion coefficient [29]. The measured rod circumference remained relatively stable below room temperature, and the impact increased with the temperature, which was managed via temperature compensation within the system (Fig. 2f, S1 Code). Together, our results demonstrate the good mechanical stability and robustness of PlantRing, which can meet the requirements of a reliable and long-term plant-wearable sensor for agricultural practices in greenhouses, orchards, and open fields.

### Versatile and high-throughput utilities of PlantRing in botanical research and agriculture

The subsequent sections demonstrate the multifaceted capabilities of PlantRing in addressing long-standing technical constraints and enhancing our understanding of plant physiology.

#### Discovering novel hydraulic mechanisms underlying fruit cracking

Fruit cracking poses a significant challenge to the quality and economic viability of crops such as tomatoes, cherries, and watermelons. Physical factors such as flesh firmness and pericarp thickness have been the primary focus in studying this trait [30, 31], whereas physiological factors including excessive rainfall or irrigation have also been linked to increased fruit cracking rate, suggesting an association between excessive sap flow into fruits and their cracking [32, 33]. However, direct evidence and a mechanistic understanding of this hydraulic phenomenon has been lacking [34]. Here, the capacity of PlantRing in measuring the sap flow distributions within the plant enabled us to tackle this daunting question (Fig. 3a). We analyzed one crack-free tomato variety (SL189) and one crack-prone variety (SL183) (Fig. 3b), together with 15 Recombinant Inbred Lines (RILs) each in four replicates derived from them that exhibited varied cracking rates. Two sensors were wrapped around the stems of each plant, one near the base and one near the truss, enabling the monitoring of water flow dynamics between the main stem and reproductive shoot at the fruit maturing stage (Fig. 3a, S6b Fig.). The acquired data revealed consistent ΔSC patterns (expansions) between the base and truss throughout the daytime (8:00–17:00) for SL189, regardless of sunny or overcast conditions (Fig. 3b). In contrast, SL183 exhibited the opposite trend, with a notable expansion in SC near the truss and a contraction at the base during the daytime, and the discrepancy is more remarkable under high VPD conditions (Fig. 3b). Similar results were observed for the crack-prone and crack-free watermelons (Fig. 3c). Since increase and decrease of SC are sensitive indicators of water influx and outflux in plant organs [35], our results suggest that the crack-prone SL183 “drew” more storage water from the main stem into its fruits, contributing to their cracking. This genotypic difference was further linked to the two varieties’ constating transpiration responses to VPD, where SL183 exhibited a more restrictive transpiration pattern at lower VPD conditions compared with SL189 (Fig. 3d), which may facilitate the maintenance and reallocation of excessive sap flow to the fruits. These findings provide novel mechanistic insights into this phenomenon from a hydraulic perspective.

**Figure 3.**
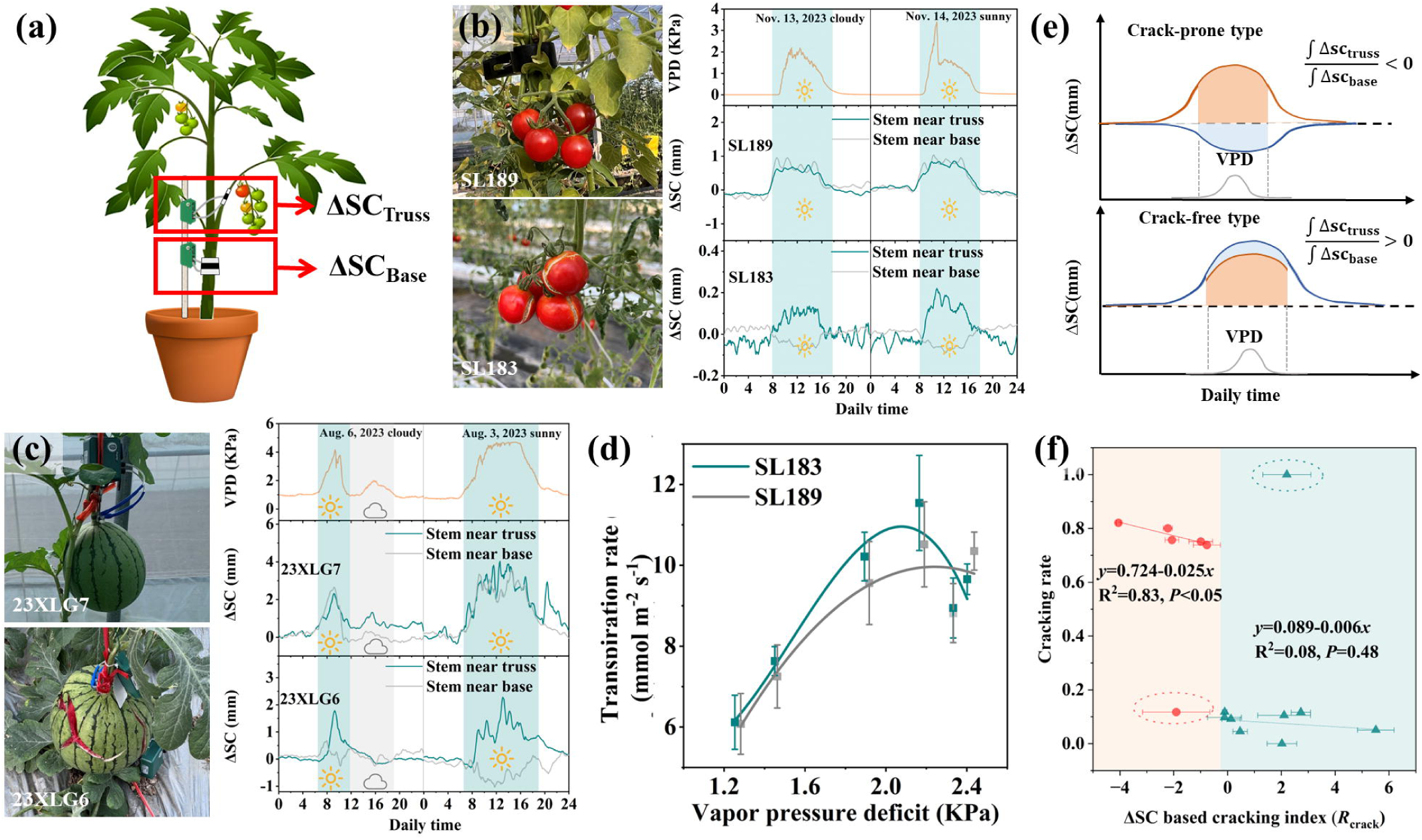
Quantified approach for assessing fruit cracking based on the measurement of stem circumference using PlantRing. (a) Principle of assessing fruit cracking using PlantRing. Two sensors are installed on each plant to measure the ΔSc of the main stem and the reproductive branch, inferring water distribution in different parts of the plant. (b) Comparison of ΔSC patterns of crack-free tomato variety (SL189) and crack-prone variety (SL183) at the base and near the truss. Data from November 13 and 14, 2023, serve as examples, with one day being overcast and the other being sunny. The ΔSC value reset to zero at 0:00 of each day. (c) Demonstration of PlantRing used to evaluate cracking index of two watermelon varieties. Comparison of ΔSC patterns of crack-free watermelon variety (23XLG7) and crack-prone variety (23XLG6). Data from August 6^th^ and 3^rd^, 2023, serve as examples, with one day being rainy and the other being sunny. ΔSC value reset to zero at 0:00 of each day. (d) Transpiration patterns of the tomato line SL189 and SL183 against daily VPD as measured by LI-600 handheld porometer (LI-COR, Inc, USA) on a sunny day (July 4, 2023). (e) Calculation of *R*_crack_, the ratio of the accumulated values of ΔSC at the base (*∫*ΔSC_base_) and near the truss (*∫*ΔSC_truss_) during the daytime. (f) Fruit cracking rates measured by independent field surveys against the cracking index (*R*_crack_) from PlantRing, using data from 15 Recombinant Inbred Lines. Data are represented as mean ± SD. Dashed circles delineate outliers. *R*_crack_ is divided into two groups based on whether they are greater than or less than 0, and each group is individually subjected to linear regression with cracking rates. Error bars represent the variation in *R_crack_* among three biological replicates. The ΔSC value reset to zero at 0:00 on December 10, 2023.

We further computed the ratio of the accumulative ΔSC at the base (*∫*ΔSC_base_) and near the truss (*∫*ΔSC_truss_) during the daytime for the RILs (offspring), which is referred to as the ΔSC-based fruit cracking index (*R*_crack_, Fig. 3e). Our data revealed significant genotypic variations in the mean *R*_crack_ during the 10-d continuous monitoring period. A strong correlation was observed between *R*_crack_ and the cracking rate measured by independent field surveys (R^2^=0.83, *P*<0.05, Fig. 3f) when *R*_crack_<0, with all these genotypes being crack-prone (cracking rates > 70%) except for one. The correlation was nonsignificant when *R*_crack_ >0, but genotypes exhibiting this characteristic also formed a distinct cluster, referred to as crack-free (cracking rate < 20%), except for one (Fig. 3f). Sporadic exceptions are expected given the complexity of the fruit-cracking trait, which is influenced by factors beyond the physiological level [34]. Therefore, *R*_crack_ can serve as an early-stage predictor of fruit cracking and as a quantitative parameter for the cracking rate in cracking-prone genotypes.

#### Quantifying stomatal sensitivity to soil drought and uncovering the novel function of a circadian clock gene

Quantifying the sensitivity of stomata to soil drought, particularly in high-throughput, has been notoriously difficult, despite being a key trait for screening drought adaptive crop lines [36]. Previous studies have established a method for measuring this trait by utilizing dynamic response curves of transpiration rate to continuously declining soil volumetric water content (VWC). In this context, the inflection point (θ_cri_) and slope (*k*) of the descending phase of the curve quantitatively define the genotype-specific responsiveness of stomata to water deficit (Fig.4a) [37]. High-throughput measurement currently relies on lysimeter arrays, such as the commercial Plantarray platform (S6a Fig.) [5, 38]. Here we demonstrate that PlantRing can perform the same function as lysimeter arrays while overcoming the need of sophisticated experimental pretreatment (e.g. repeated water saturation of the medium) and high instrumentation cost. Fig. 4a shows a comparison of using Plantarray and PlantRing over 10 days on identical tomato plants. During the initial days of progressive soil drought before reaching a particular VWC threshold (θ_cri_), the midday (12:00–14:00) whole-plant transpiration normalized to VPD (Tr_mid,VPD_) measured by Plantarray remained rather constant, whereas midday stem circumference relative to the initial value (ΔSC_mid_) measured by PlantRing showed a slight linear increase, reflecting the continuous daily stem growth. Importantly, the two systems detected very similar VWC thresholds (Fig. 4a). Beyond these thresholds, both Tr_mid,VPD_ and ΔSC_mid_ exhibited a linear decrease in relation to declining VWC at a similar rate. Therefore, PlantRing offers an alternative approach for detecting θ_cri_ and *k* with minimal experimental requirements. We subsequently applied this approach to a collection of common bean varieties, a globally essential legume crop. By simultaneously monitoring the dynamics of ΔSC_mid_ in different genotypes undergoing progressive soil drought, we were able to distinguish sensitive (θ_cri_>0.23) germplasm from insensitive (θ_cri_<0.17) germplasm based on the accurately quantified θ_cri_ and *k* values (Fig. 4b). This proves the potential of PlantRing for screening drought-adaptive lines from large germplasm collections.

**Figure 4.**
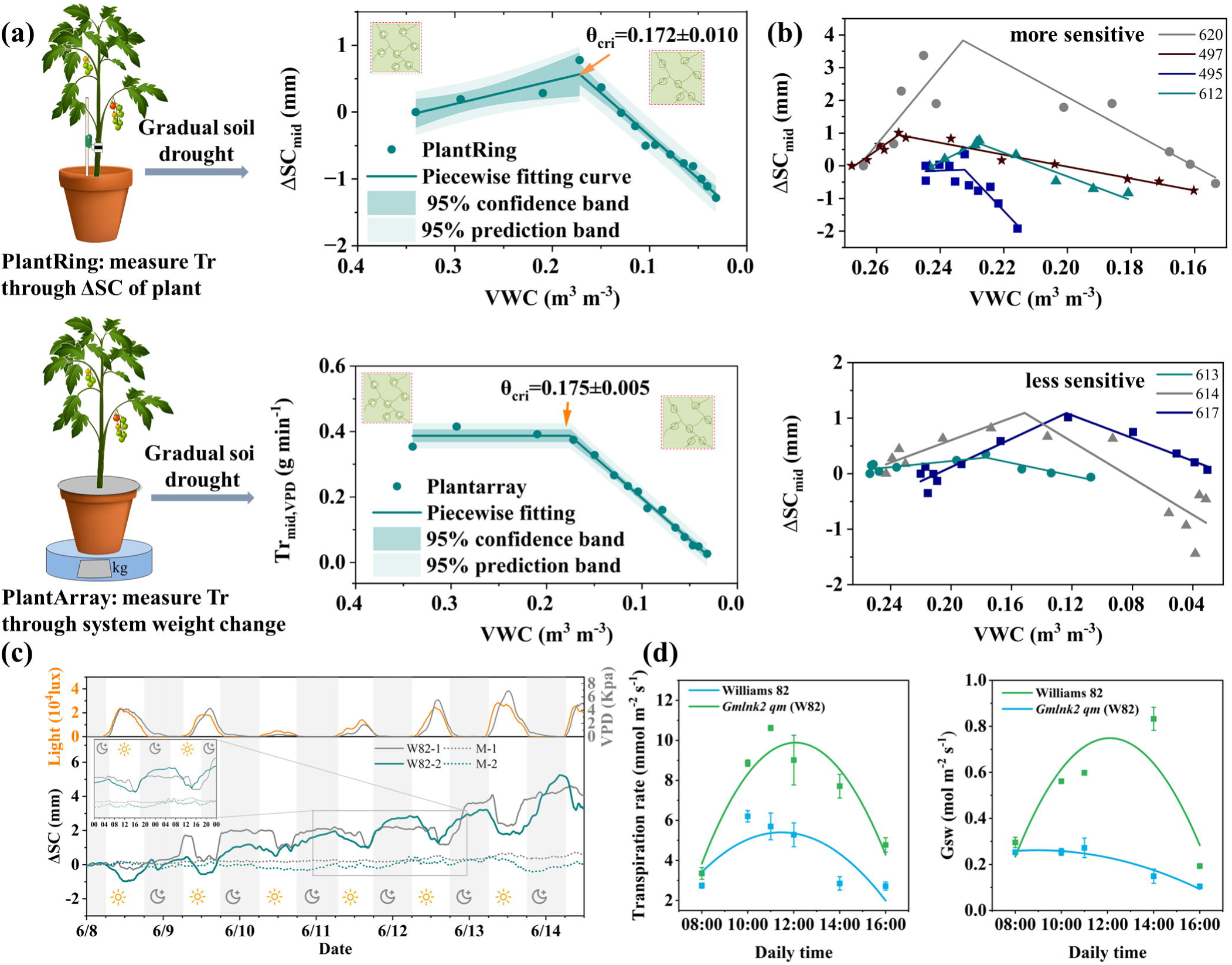
Detecting stomata sensitivity to soil drought in various crops and the diurnal ΔSC in wild-type and *Gmlnk2 qm* mutant soybean plants. (a) The principles of measurement using the two different systems. Plantarray and PlantRing measure transpiration level through system weight variation and variation of stem circumference, respectively, both of which are the results of water uptake/loss from the plants. The midday (12:00–14:00) stem circumference (ΔSC_mid_) or transpiration of tomato plants normalized to VPD (Tr_mid,VPD_) are fitted with relative soil VWC. The response curves are fitted using a two-piecewise function, where θ_cri_ represents the inflection point at which ΔSC_mid_ or Tr_mid,VPD_ significantly decreases with VWC. (b) Measurement with PlantRing classified seven common bean genotypes into the sensitive and insensitive groups in terms of stomatal response to gradual soil drought, based on the θ_cri_ values. For clarity, data of one representative plant each are presented. (c) Comparison of the ΔSC patterns between wild-type soybean (W82) and its quadruple mutant (M) on the *GmLNK2* gene from June 8 to June 15, along with corresponding changes in light intensity and VPD throughout the measurement period. For clarity, data of two representative plants each are presented. Data from June 11 to June 13 are highlighted to show the diurnal variations in ΔSC. The ΔSC value was reset to zero at 00:00 on June 8. (d) Transpiration rate (Tr) and stomatal conductance (Gs) patterns measured by LI-600 handheld porometer on a sunny day.

Circadian clock genes are crucial for plant functionality, and certain of these genes have been associated with more diverse functions [39]. We utilized PlantRing to investigate the SDV patterns of the wild-type soybean Williams 82 (W82) and its quadruple mutant (M) for the gene *NIGHT LIGHT–INDUCIBLE AND CLOCK-REGULATED 2* (*LNK2*) with a known function in circadian period and flowering time control (Fig. 4c) [40]. In W82, measurement of SDV showed daytime contraction and nighttime expansion, which aligned with a transpiration-related sap flow model. In contrast, the mutant exhibited a profoundly diminished diurnal fluctuation in SDV, indicating impaired stem sap flow distribution responding to the environment due to the loss of *GmLNK2* function. Manual measurements further revealed a significantly reduced transpiration rate (Tr) and stomatal conductance (Gs) level in the mutant throughout the day, along with a more pronounced loss of diurnal Gs fluctuation (Fig. 4d). Overall, our system provides a robust method for measuring the biological rhythm of sap flow via SDV and suggests a novel function of *GmLNK2* in stomatal regulation, which may or may not relate to its known function.

#### Plant-based feedback irrigation empowered by PlantRing

As PlantRing offers real-time monitoring of water status through ΔSC, we developed an automated feedback irrigation system incorporating its function (Fig. 5a). This irrigation system allows access and response to plant-based information to govern irrigation instead of relying on user experience or environmental parameters such as VWC. In a real agricultural tunnel experiment with tomato plants, three irrigation modes were implemented (Fig. 5b-d): well irrigation (WI), deficit irrigation based on soil VWC (DI_v_) and deficit irrigation based on PlantRing (DI_p_). During the 31-day experimental period, the total water consumption in the DI_p_ treatment, calculated based on irrigation duration and water discharge rate, was approximately 1/2 and 1/3 of that for the WI and DI_v_ treatments, respectively. The fruit fresh weight per plant of DI_p_ group was slightly only lower than those of WI and DI_v_ groups, whereas the dry weight showed no significant differences (Fig. 5e). More intriguingly, the soluble solid content, a key quality trait, was 14.7% and 6.3 % higher in the DI_p_ group compared to the WI and DI_v_ groups, respectively (Fig. 5f). In a similar experiment conducted under artificial growth chamber condition, the DI_p_ group demonstrated simultaneous water saving and improved fruit quality with no yield loss compared to the WI group (S7 Fig.). These results demonstrate the promising utility of PlantRing for guiding plant-based deficit irrigation as a next-generation approach.

**Figure 5.**
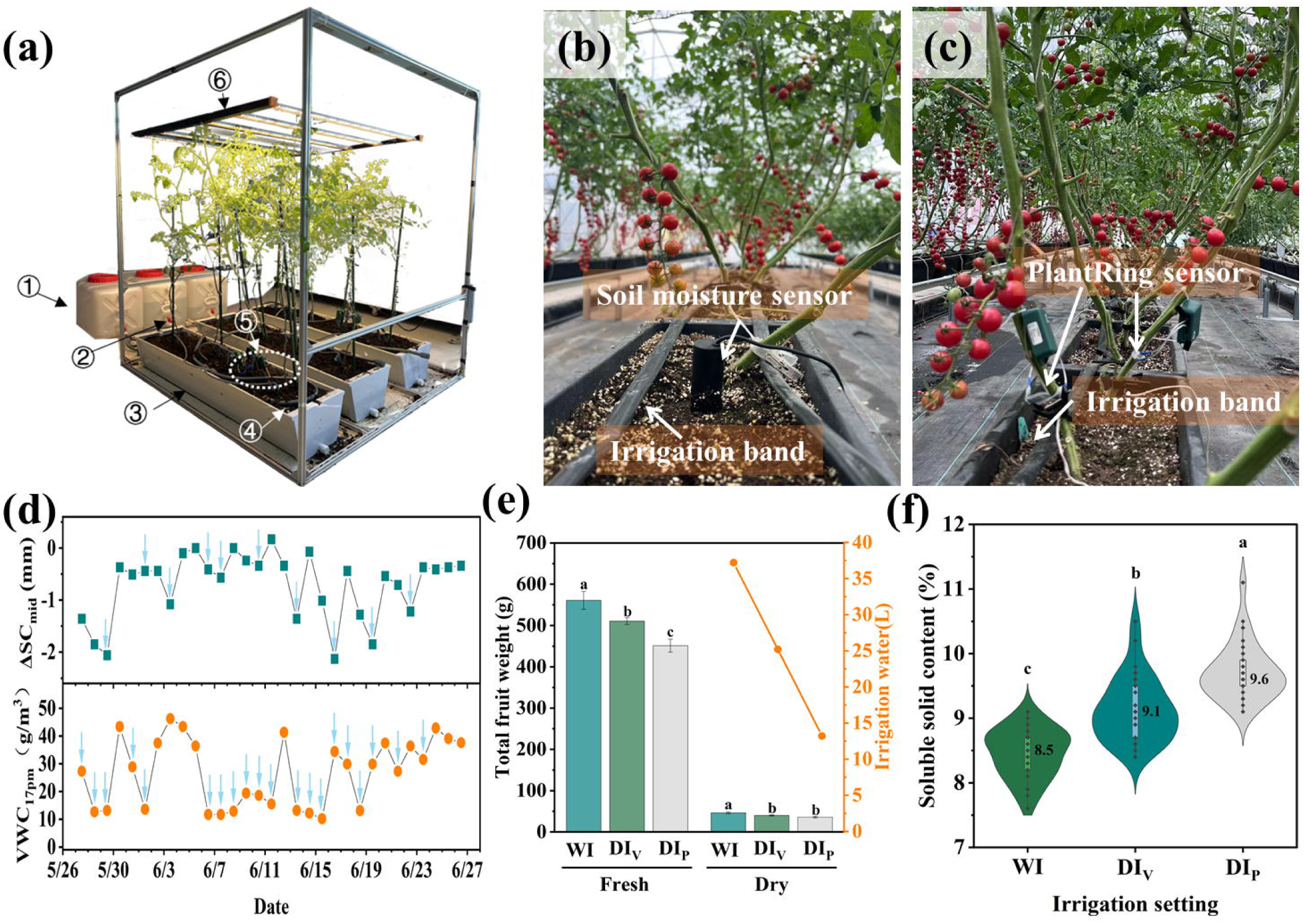
Plant-based feedback irrigation empowered by PlantRing. (a) Prototype design of PlantRing-based feedback system. The plants were cultivated in trapezoidal PVC plant cultivation troughs with an outlet ③ and full spectrum LED lights ⑥ were employed to provide illumination, a mini-pump ② was used to extract water from tank ① through irrigation pipe ④ and irrigate the cultivation troughs according to the feedback of PlantRing systems installed on the plants ⑤. (b) Real-experiment image showing traditional feedback irrigation system based on soil moisture sensor and automatic irrigation system with irrigation band. (c) Real-experiment image showing feedback irrigation system incorporating PlantRing. (d) The data record for irrigations as feedback based on soil moisture sensor and PlantRing. Less irrigation was achieved when using PlantRing. (e) Fresh and dry weights of all fruits per plant in well irrigation (WI), deficit irrigation guided by soil VWC (DI_v)_, and deficit irrigation guided by PlantRing (DI_p_) treatments, with four plants included in each treatment group. Data are represented as mean ±SD. Different lowercase letters and “*” denote significant differences between treatments (*P*<0.05), utilizing the pairwise comparison method of least significant difference (LSD). The orange line demonstrates the total amount of irrigation solution used for the three treatments during the experiment. (f) Soluble solids content of fruits in WI, DI_v_, and DI_p_ treatments, with four plants included in each treatment and six fully ripe fruits in each plant, were selected. The maximum value (top of plot), 75th percentile (top of box), 50th percentile (median), 25th percentile (bottom of box), minimum value (bottom of plot) and outlier were shown from top to bottom in the Violin-plot. The scatter points represent the raw data, the average value is numerically presented on the right side of box.

## Discussion

The following decade is expected to undergo a transition in plant wearables from conceptual or theoretical demonstrations to large-scale applications. PlantRing features a lightweight design surpassing LVDT sensors and integrates multiple advantages, simultaneously achieving ultrahigh sensitivity and substantial stretchability, along with notable robustness, durability and stability under agricultural conditions. To our knowledge, PlantRing is currently the only device with proven versatile use for high-throughput monitoring the dynamic organ growth and water relations under real agricultural conditions. Unlike thermal transport-based wearable sensors, which are limited to herbaceous plants and have a restricted sensing range [13, 14], PlantRing is widely applicable to various plants and its functionality can be extended from agriculture to forestry by measuring the dynamics of diameter at breast height, a key parameter in woody species that reflects their growth and health status [41]. This is intended to introduce a technical revolution in forestry by replacing traditional manual measurements and rigid LVDT sensors. In addition, the sensor fabrication process of PlantRing is easily standardized, and the sensors can be treated as disposable in applications owing to their convenient plug-out design and low cost. The multi-modular design allows for separate pre-production and assembly, offering more flexibility for commercialization compared to on-site fabrication and printing technologies [42].

Implemented on a large scale, PlantRing has proven invaluable in understanding genotypic variations in the growth and physiological traits. We demonstrate the use of PlantRing to trace dynamic stomatal behaviors as a function of ΔSC during gradual soil water depletion, and to quantify the critical points (θ_cri_) and rates (*k*) of stomatal closure. This addresses a long-term challenge in drought studies. The ability to acquire population-level θ_cri_ and *k* data will enable forward genetic mapping of complex stomatal behavioral traits, leading to the identification of genes/QTLs governing them. PlantRing demonstrated another advanced utility by elucidating the hydraulic mechanism behind complex fruit-cracking traits. In traditional visual phenotyping, fruit cracking often appears as a late symptom and is frequently unstable [43]. Early-stage measurements using texture analyzers are destructive and not conducive to efficient high-throughput phenotyping [44].Our findings reveal distinct ΔSC patterns between main stem and fruiting branch in crack-prone versus crack-free varieties, raising the theory that asynchronous intra-plant sap flow distribution related to transpiration patterns underlies fruit cracking. This introduces a new factor influencing fruit cracking [34]. We also established and validated R_crack_, based on cumulative ΔSC, as a physiological predictor for fruit cracking. Continuous monitoring of ΔSC patterns can guide timely interventions, such as soil moisture management, to reduce fruit cracking rates. Additionally, our findings suggest that modifying or regulating plant transpiration responses to VPD may offer a potential strategy for mitigating fruit cracking. Furthermore, the observation of a daytime increase of stem circumference in both main stem and fruiting branches in the cracking-free tomato and watermelon genotypes is noteworthy, highlighting that the sap flow direction in plants are complex and dynamic traits not only related to diurnal environmental change but also to plant developmental stage and genotype [45, 46] Combing PlantRing with more physiological and molecular tools will provide a deeper understanding of the phenomenon.

Incorporating PlantRing into feedback irrigation marks a paradigm shift in smart agriculture, moving from an experience- or environmental parameter-driven approach to a direct plant information-driven mode. Traditional feedback irrigation relies on monitoring soil moisture, but this may not necessarily reflect the plant water status [42]. Existing conceptual plant-based monitoring techniques for feedback irrigation include monitoring leaf turgor pressure, leaf thickness, sap flow and xylem cavitation [47]. However, owing to the considerations of cost, operational simplicity, and the intricate complexities of data parsing, their utilization in commercial applications has been limited [48]. PlantRing, with its cost-effectiveness and well-established algorithms linking measurements to plant water status, has the potential to become a game changer. The key to a deficit irrigation schedule is identifying the appropriate point for initiating irrigation in repeated wet-dry cycles [49]. Using PlantRing, we found the optimal deficit irrigation threshold by analyzing the relationship between daily minimum ΔSC and maximum Tr for three days. This approach resulted in increased fruit sugar content with less watering (∼1/3) and minimal decrease in yield, possibly due to reduced fertilizer input with irrigation, compared to well-irrigated crops. In practical applications, the length of the day window for calculating the deficit irrigation threshold need to be optimized according to specific crop types and soil or substrate properties. Compared with the approaches based on indirect environmental parameters such as VWC, our ΔSC-directed feedback irrigation reduces the need to quantify the environment-plant correlation through system modeling strategies, which is typically the most time-consuming and imprecise aspect in practice [50].

PlantRing has substantial potential for various other versatile applications. For example, future research could explore its utility in dissecting complex traits, such as die-off points to abiotic stresses, by integrating SC measurements with mechanistic models [18, 28, 51]. The capability of PlantRing to unveil the relationship between nutrient status and organ circumference also warrants further exploration and holds the potential to guide intelligent fertilization practices. Such capacity, combined with genetic and mutant analysis as demonstrated in this study, will boost identification of key genes responsible for the traits. Potential limitations of PlantRing include, first, the challenge of distinguishing between irreversible growth effects and reversible hydraulic effects when measuring rapidly growing organs. However, both our data and previous study indicate that, on a daily basis, the impact of the former on SDV is typically minor compared to the latter. Secondly, the resolution of PlantRing is currently constrained by the resistance measuring module in the data logger, a compromise made to balance performance and cost. However, this resolution can be enhanced to a higher level of 10 μm with more advanced electrical testing instruments. Further enhancements of the PlantRing system could also involve using an autonomous energy harvester powered by plants, rain, or wind. Harnessing advancements in deep learning technologies through data training will allow accurate estimation of growth and water-related parameters from SDV datasets [52]. The combination of PlantRing with other wearable sensors may expand its functionality to a more comprehensive level.

## Materials and methods

### Sensor fabrication

The sensor fabrication protocol was developed based on the preparation procedure of carbonized silk georgette (CSG)-based strain sensor reported in our previous work [24]. Briefly, a commercial silk georgette was carbonized at high temperatures under an inert atmosphere (continuous gas flow of 100 sccm argon and 10 sccm hydrogen). The heat treatment consisted of four stages, as presented in S2 Table. Rectangular strips of CSG were then prepared to the required length for the strain sensor. A silver adhesive was applied to both ends of the strip to connect it to a piece of flexible printed circuit connector for electrical signal measurement. The CSG strip was placed on a solidified polydimethylsiloxane (PDMS) substrate with an approximate thickness of 450 μm. The PDMS substrate was formed by blending the base and cross-linker at a weight ratio of 10:1, removing air from the mixture, and allowing it to solidify at 80 °C for 3 h. The liquid PDMS precursor was then coated onto the CSG and left to dry. The sensor component, which was enveloped in latex, was designed to be waterproof.

### System integration

The interconnection between the strain sensors and data logger was achieved through a sophisticated assembly comprising flexible printed circuit connectors, wires, and banana plugs and sockets (Fig. 1a). The data logger collected resistance change of the sensor and transmitted the signal wirelessly to a gateway (Fig. 1d, S2 Fig.). The gateway offered the option of inserting an Internet of Things (IoT) card or connecting using an Ethernet cable, enabling communication with external networks through 4G/5G/WLAN technologies for seamless interaction with servers (Fig. 1e). The PlantRing cloud-monitoring platform (112.124.0.216:8080), compiled in Java, was hosted on Alibaba Cloud servers to guarantee reliable accessibility. The functionalities of the cloud platform included a homepage display with system usage instructions, gateway management, management of data transmission terminals, monitoring object management, experiment management, and system administration (S3 Fig.).

### System operation and calibration

The PlantRing system was operated remotely via a cloud platform. Gateways were placed in protected or open fields with access to power and the internet to facilitate the connection of data loggers. Each data logger was matched to an assigned strain sensor unit. The signal was calibrated with a uniaxial measurement device (S4 Fig.) to guarantee consistent results across different sensor batches. The sensor’s two ends were fixed with pliers on the device, and the strain level was precisely adjusted by moving one of the pliers along the sliding rail. The variation in resistance was recorded and plotted against the stretched length of sensor (Fig. 2a). The slope of plot was programed onto the PCB via a type-C connector to complete the sensor calibration process. The sensor was connected to a data logger with a banana plug inserted into the corresponding socket. The cloud platform allowed users to configure the desired frequencies for data collection (default: 3 min; minimum: 1 sec) and intervals for gateway reporting. Using the monitoring panel interface, users can select specific sensor datasets and define custom ranges for graphical representation and subsequent export.

### Evaluation of the system performance

The resolution of PlantRing was determined by the smallest distance at which the ADC signal began to change. Repeatability was assessed by stretching the sensor to the same strain level and calculating the coefficient of variation. Accuracy was calculated by comparing the ratio between the stretching distance deducted from ADC signal value and the actual stretching distance.

The system stability under wind or rain interference was evaluated using 45-day-old tomato plants (cv. Alisa) in 2 L pots filled with a nutrient-rich soil substrate (Zhonghe Co., Ltd., China). Each experimental group consisted of three (wind interference) or four pot (rain interference), each housing a single tomato plant equipped with a PlantRing attached to its stem base. To assess the impact of wind, plants were placed in a tunnel with natural light and ambient temperatures (ranging from 25 to 34°C on the testing day, November 28th, 2023). An artificial airflow at a speed of approximately 3.5 m/s was generated by an oscillating fan (AD61-1, AIRMATE, China) positioned 1 m away from the tomato plants (S1 Movie). December 4, 2023, which was a rainy day, was chosen for the rain impact assessment. The control pots were kept indoors, whereas the treatment group pots were placed in an open field (S2 Movie).

To compensate for the impact of temperature, ten sensors were attached to a quartz glass rod with a diameter of 7 mm, which has a thermal expansion coefficient close to zero. The rod was placed in a DHG-9423A oven (Jinghong Experimental Equipment Co., Ltd., China). The test began at 5 °C, with the temperature gradually increasing by 5 °C every 10 min until reaching 50 °C. Sensor readings (denoted as x) were collected every 3 mins and plotted against time (denoted as y) to establish the correlation between sensor readings and temperature changes. The measured length of the sensor was adjusted to match the value obtained at room temperature, using temperature data from the sensor embedded in the PCB of the data logger. The general principle included randomly selecting 5 sets of data from the 10 tested sensors and using the SimpleImputer tool to fill in the missing data. We then calculated and compared the mean squared error of a linear regression model and a quadratic polynomial regression model to determine the optimal method. The entire procedure was executed using S1 Code provided in Support Information. The sensor length could be compensated using the following equation: *y* = - 0.03362482 *x* + 0.00062241 *x*^2^.

### Measurement of organ circumference using PlantRing

Tomato (cv. Alisa) plants were cultivated in 2 L pots filled with a nutrient-rich soil substrate (Zhonghe Co., Ltd., China). Cultivation occurred in a glass greenhouse with natural light from May 6^th^ to May 13^th^, 2024 and meticulous temperature control, maintaining a daytime range between 20 and 35 °C. When the FC reached approximately 140 mm, the PlantRing sensors were applied to the tomato fruits to monitor the dynamic changes in FC continuously. Additionally, every two days at 9:00, we manually measured the fruit diameter (FD) using the DL91150 digital caliper (Delixi Group Co., Ltd., China) and calculated the FC using the formula FC=πFD.

Soybean (*Glycine max*) materials used in this study include the wild-type variety Williams 82 (W82) and the *Gmlnk2 qm* mutant [53]. Each plant was cultivated in 2 L pots filled with a nutrient-rich soil substrate (Zhonghe Co., Ltd., China). The cultivation took place in a glass greenhouse with natural light and temperature control in May to June 2024, maintaining daytime temperature between 30 and 45 °C. Forty days after seed sowing, a PlantRing sensor was attached to each plant at the stem base to monitor dynamic variations in SC.

### Measurement of stem diameter using laser displacement sensors

High-sensitive laser displacement sensors (HL-T1010A with HL-AC1 controller, Panasonic Corporation, Japan) with an 8 μm resolution were deployed in parallel with PlantRing to provide a comparison of the dynamics of ΔSC, based on the measured stem diameter (SD) (S5 Fig.). To prevent swaying during measurements, the tomato plants were securely held in place using supporting frames (S5b Fig.). The experiment involved three distinct stages: well irrigation (WI), progressive water deficit (WD), and rehydration (WR). In the WI and WR phases, the nutrient solution was provided by drip irrigation for 240 s (oversaturated) at 23:00, 1:00, 2:00, and 3:00 of the day. No nutrient solution was supplied during the WD stage.

### Phenotyping of the common beans using PlantRing

Seven varieties of common bean (*Phaseolus vulgaris* L.) were used. Each plant was cultivated in 1.5 L pots, all containing precisely measured equal amounts (1.5 kg) of soil enriched with a nutrient-rich substrate (Pindstrup Mosebrug A/S, Ryomgaard, Denmark). Three biological replicates were set for each variety. Cultivation occurred in a glass greenhouse with natural light and temperature control in June 2024, maintaining a daily temperature between 25 and 38 °C. Sixteen days after seed sowing, when the plants reached the mature stage with minimal growth effects on SC, a PlantRing sensor was attached to each plant’s stem base to monitor dynamic variations in SC. Gradual soil drought was initiated by withholding irrigation on the same day that the sensors were mounted. Soil VWC in each pot was measured using the 5TM soil sensors.

### Phenotyping fruit cracking potential for tomatoes and watermelons using PlantRing

Fifteen F_6:8_ RILs, resulting from a cross between SL183 (crack-prone parent) and SL189 (crack-free parent), along with the parental lines, underwent simultaneous phenotyping using PlantRing between September and December 2023 and were each assessed in four biological replicates. The plants were cultivated in troughs filled with nutrient-rich soil substrate within a glass greenhouse with natural light and meticulous temperature control, maintaining a daytime temperature range between 20 and 35 °C. Regular automatic weekly irrigation with drippers was conducted to maintain the soil moisture. Key milestones included transplanting on October 7, trellising on October 25, the initiation of flowering on November 10, and the fruit onset on November 20. During the tomato extension phase, the lateral branches were pruned, and only one truss was retained. On December 10, two PlantRing sensors were affixed to each plant: one each near the stem base and the truss.

### Phenotyping using a lysimetric array (Plantarray)

To facilitate a comprehensive comparison with the PlantRing, physiological phenotyping, including the assessment of transpiration and stomatal sensitivity, was also conducted using the commercial high-throughput physiological phenotyping system Plantarray 3.0 (Plant-DiTech, Israel). The Plantarray assay followed established procedures [54]. Briefly, 25-day seedlings were transferred to the load cell of the plant array. To prevent the evaporation of soil water, the pot surface was wrapped with a plastic film. Soil VWC was obtained from the 5TM soil sensors integrated into the system, and environmental VPD was calculated from relative humidity (RH%) and air temperature (℃) acquired by the system’s sensor. The whole-plant transpiration at the 3-min step was calculated by multiplying the first derivative of the measured load-cell time series by -1 [5]. Daily whole-plant midday transpiration (Tr_m_, averaged over the period between 12:00 and 14:00) was fitted to a piecewise linear function of the corresponding VWC during the dynamic period of water deficit [55]. To offset the influence of daily environmental variations, Tr_m_ was normalized to VPD (Tr_m,VPD_).

### Fabrication of the prototype plant-based feedback irrigation system

The prototype feedback irrigation system incorporating PlantRing was fabricated comprising horizontal plastic water tanks (dimensions: 70 × 33 × 33 cm), trapezoidal PVC plant cultivation troughs with an outlet (length: 120 cm, height: 18 cm, top width: 30 cm, bottom width: 20 cm), a full-spectrum, dimmable LED plant growth light (Yihao Agricultural Technology Co., Ltd, China), a 60 W miniature diaphragm pump (Pulandi Mechanical Equipment Co., Ltd, China), irrigation main pipes (outer diameter: 16 mm, inner diameter: 13 mm), an adjustable nozzle (Zeego, China), the PlantRing sensors, and an agricultural IoT system (Yihao Agricultural Technology Co., Ltd, China) for comprehensive control. The port of the PlantRing cloud platform was accessible for integration with the aforementioned agricultural IoT system. Consequently, the SDV data of plants could be seamlessly transmitted to the IoT system, enabling the initiation of different irrigation programs.

### Settings of the feedback irrigation experiments

‘Micro Tom’ tomato plants were grown in cultivation troughs filled with nutrient-rich soil substrates. The experiments were conducted both in an artificial growth chamber environment with the aforementioned prototype feedback irrigation system and in a glass tunnel for real production, following similar principles.

For the artificial growth chamber experiment, the daily temperature range was controlled between 20 and 30 °C. Light intensity simulated natural daily variations, starting at 20% at 7:00 and increasing to 50% at 9:00, further increasing to 80% at 10:00, peaking at its maximum Photosynthetic Photon Flux Density (PPFD) of 1720 μmol s^-1^ at noon. In the afternoon, it gradually decreased to 60%, 40%, and 20% at 15:00, 16:00, and 17:00, respectively, finally setting at 0% at 19:00. Irrigation was performed every alternate day during the WI treatment. In the SD treatment, watering was initiated by the empirical observation of leaf wilting. As for the DI treatment, irrigation was determined based on ΔSC. The ΔSC_mid_ over five consecutive days (including the current day and the preceding four days) was linearly fitted, and irrigation commenced when the slope was less than 0 (Fig. 5d). In all treatments, the watering phase occurred at 8:00, lasting for 60 s to ensure adequate hydration, with each plant equipped with one drip head and an approximate water discharge rate of 600 mL/min per drip. The experiment concluded when approximately half of the fruits on all plants had ripened.

The tunnel experiment was performed during May to June 2024, with natural light and controlled temperature between 20 and 35°C throughout the day. The irrigation of three experiment groups was controlled through an integration system of irrigation. The cultivation troughs of each group shared an irrigation band connected to a faucet, which could release a fertilizer solution into the soil. A solenoid valve inside the faucet controlled the timing and volume of irrigation, and was remotely operated with a key station that automatically programmed the irrigation plan based on the feedback from PlantRing or soil moisture sensors. The irrigation plan was as follows: (1) WI group: irrigated every day; (2) DI_v_ group: irrigated if the VWC measured by soil moisture sensor (YHW02-2, Yihao Agricultural Technology Co., Ltd, China) was lower than 75%, according to the protocol by Huffman et al. [56]; (3) DI_p_ group: irrigated if linear regression of the minimum ΔSC values recorded until 17:55 during the daytime shows a negative slope over three consecutive days. Once the irrigation requirement is met, 1.2 L of fertilizer solution was pumped into the cultivation trough at 18:00 for all three groups.

### Measurement of fruit weight and soluble solids

When tomatoes reached the ripening stage, four plants were selected from each treatment, and the number and fresh weight of fruits were measured for each plant. Subsequently, six fully ripened fruits from each pot were selected for testing soluble solids. The fruits were crushed, their uniform pulp was carefully placed into the measuring hole of a PAL-1 pocket refractometer (ATAGO, Japan), and accurate readings were recorded. After each measurement, thorough cleaning was performed using pure water. After the tests, all fruits were dried to a constant weight in a DHG-9423A oven (Jinghong Experimental Equipment Co., Ltd., China) at 80 °C, and the dry weight was measured.

## Supporting information

Support information

## Acknowledgments

We express our gratitude to Dr. Xia Cui from the Chinese Academy of Agricultural Sciences for generously providing us with the populations of tomato RILs.

## Support Information

**S1 Fig. Diagram of the clip design for sensor adjustment and fixation.** (a) The clip on the sensor can be opened to adjust the length required for testing. (b) Once the desired length is achieved, the clip can be locked in place by snapping it shut. (c) Image showing a PlantRing system installed on a tomato stem.

**S2 Fig. Overview of the data logger.** The sensor unit comprises the following components: ① a strain sensor, ② wires with banana plugs, ③ banana sockets, ④ a Type-C interface, ⑤ a power/zeroing button, and ⑥ an LED indicator light. The data logger is rated IPX5 for water resistance, protecting against water projected from a spray nozzle at any angle for 10 to 15 minutes from a distance of 3 meters at a pressure of 30 kPa. The PCB integrates the NRF52833 Bluetooth chip and an ADC module, powered by a dedicated charging management system connected to a 380 mAh battery. It includes operational amplifier circuitry for connecting strain sensors and an air temperature and humidity sensor via an Inter-Integrated Circuit (I2C) protocol. User interaction is facilitated through General-Purpose Input/Output (GPIO) connections, which support an LED light and a multifunction button, enabling operation control, resetting, or shutting down the device. The PCB design also provides for easy battery charging and serial data download via a Type-C interface. Additionally, the PCB is equipped with a PCB antenna, and during operation, the data logger communicates with the gateway using 2.4G RF technology.

**S3 Fig. Overview of the cloud server interface.** (a) Homepage Layout: The homepage showcases functionalities such as language selection (Chinese and English), full-screen mode, and webpage locking. An example displays shows temperature, humidity, and length values. (b) Gateway Status and Settings: The network currently encompasses over forty gateways, each capable of managing at least 300 PlantRing units. The reporting and sampling intervals for these gateways can be adjusted, as highlighted on the right side of the figure. (c) Sensor Information in Experimental Design: Groups can be created based on experimental parameters to review data from different gateways. The example illustrates how to search and access specific sensor information, including plant species, detection target, sensor ID, and historical data. In this instance, the experimental group is labeled "Control group" with tomato as the plant species, stem as the detection target, and "38FBD30F70B1" as the sensor ID. (d) Data Export for Specific Date Ranges: Users can select a specific date within the timeline to view the 24-hour length curve for that day. By clicking the "Export" button, an Excel file containing time, length values, temperature, humidity data, sensor ID, and battery level can be downloaded. The four icons arranged from left to right offer functionalities for data visualization, line graph representation, bar chart depiction, and curve image preservation. Users can select a start and end date within the timeline, and clicking "Confirm" will display the curve for the chosen date range. The "Smooth" button applies a 20-point smoothing algorithm to the curve, and the "Reset" button allows users to select a new date range.

**S4 Fig. The uniaxial measurement device used for signal calibration of sensors.** The device consists of two pliers, one installed on a fixed platform and the other on a mobile platform. The sensor was secured at both ends by the pliers, maintaining its original length at the starting point. The mobile platform can be moved with precise distance along the slide rail to control the strain applied to the sensor. A spacer could be inserted to keep the distance constant. During the stretching process, the sensor was connected to the data logger and the variation of resistance was recorded on the cloud sever. The resistance value was then plotted against the stretched length of the sensor, with the slope of the plot used for signal calibration.

**S5 Fig. Measurement of stem diameter using a laser displacement sensor.** A laser displacement sensor was positioned near the probes at both ends of the PlantRing that was attached to the stem. These sensors were aligned using a custom-designed linear track, ensuring that the stems of the tomato plants consistently remained within the laser beam’s range. The light spot was partially occluded by the stem, which could be used to measure the stem diameter with the receiver end.

**S6 Fig. Images of plants grown Plantarray and/or equipped with PlantRing.** (a) Common bean plants were cultivated on the Plantarray lysimeter platform and installed with the PlantRing system to compare measurement of water relations. The Plantarray platform dynamically measures the total weight of system, whereas PlantRing system dynamically measures ΔSC of the main stem. (b) Tomato plants each installed with two PlantRing sensors for studying fruit cracking based on the measurement of stem circumferences. One sensor was installed on the main stem, and the other on the fruiting branch.

**S7 Fig. Results of PlantRing-based feedback irrigation under laboratory artificial lighting conditions.** The results demonstrate the promising utility of PlantRing for guiding plant-based deficit irrigation as a next-generation approach to achieve water conservation and quality improvement simultaneously. (a) Three irrigation modes were implemented for cultivation of tomato plants, the ΔSC was recorded for comparison. The three modes included: (1) regular irrigation every 2 days (well irrigation treatment, WI), (2) irrigation initiated upon observation of leaf wilting (severe drought treatment, SD), and (3) irrigation initiated when a negative slope was detected using linear regression for the ΔSC_mid_ over five consecutive days, indicating a significant decrease in ΔSC_mid_ (deficit irrigation treatment, DI). (b) Fruit productivity under different treatment, measured by the fresh or dry weight of all fruits per plant. The results showed no significant difference between the WI and DI groups, whereas a reduction in yield was noted under SD treatment. A reduced number of fruits per plant (by 26.7%, P<0.01), rather than reduced fresh or dry weight of individual fruits (P>0.05), was the reason for yield reduction in SD treatment. (c) The measured soluble solid content of fruits, which showed a notable increase in DI compared with WI and SD.

**S1 Table. The heating procedure for producing carbonized silk georgette.** The temperature was adjusted from the starting temperature to the heating temperature at different rate, then the temperature was kept constant for different heating time before the next stage starts.

**S2 Table. Characterization of the repeatability and accuracy of PlantRing system.** The data was obtained from testing 9 different 6 cm type sensor, AVD AD stands for the average AD value measured under different levels of strain, STD stands for the standard derivation of the AD values, CV stands for the coefficient of variation of the AD values, MS stands for measured strain deducted from AD value, which is also displayed and recorded in cloud sever, RE stands for the relative error calculated by comparing measured strain with the actual strain.

**S1 Movie.** The experimental scenario for testing PlantRing under controlled airflow conditions. The tomato plants were placed in a tunnel with natural light and ambient temperatures (ranging from 25 to 34°C on the testing day, November 28^th^, 2023). Three tomato plants were selected for the experiment, each equipped with a PlantRing attached at its stem base. An oscillating fan (AD61-1, AIRMATE, China) was positioned one meter away from the tomato plants to generate an airflow velocity of approximately 3.5m/s. The wind interference period lasted from 9:00 to 16:00.

**S2 Movie.** The experimental scenario for testing PlantRing under rainy day. The tomato plants were placed in an open field in a rainy day (December 4^th^, 2023). Four tomato plants were selected for the experiment, each equipped with a PlantRing attached at its stem base. The rain interference period lasted from 10:00 to 16:00, during which the precipitation rate ranged from 0.1 to 0.38 mm/h.

**S1 Code. The code used for temperature compensation.**

The measured rod circumference remained relatively stable below room temperature, and the impact increased with the temperature, which was managed via temperature compensation within the system. The code for temperature compensation is as follows:

