## Supplementary material for "PlantRing: A high-throughput wearable sensor system for decoding plant growth, water relations and innovating irrigation": Support information

**This PDF file includes:**

Materials and methods
Figures S1 to S7
Tables S1 to S2
Legends for Movies S1 to S2
Code S1
SI References

**Other supporting materials for this manuscript include the following:**

Movies S1 to S2

(a) (b) (c)

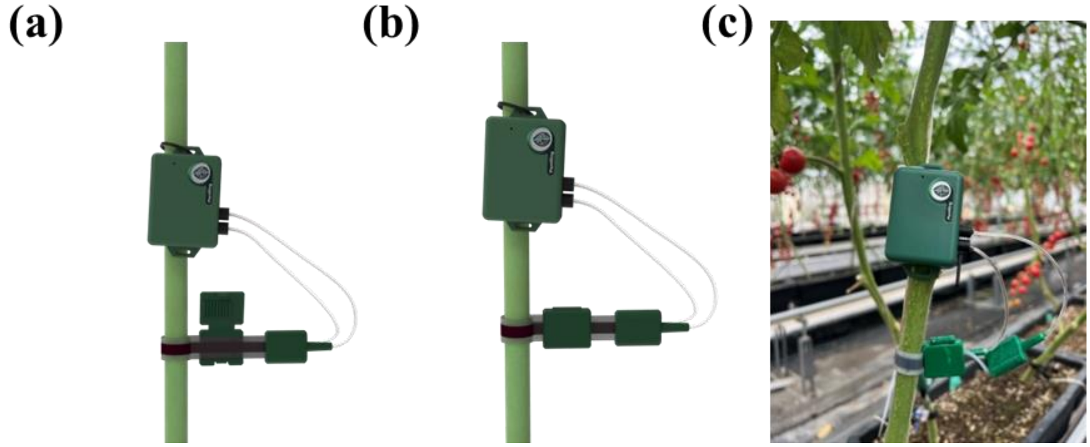

**Fig. S1. Diagram of the clip design for sensor adjustment and fixation.** (a) The clip on the sensor can be opened to adjust the length required for testing. (b) Once the desired length is achieved, the clip can be locked in place by snapping it shut. (c) Image showing a PlantRing system installed on a tomato stem.

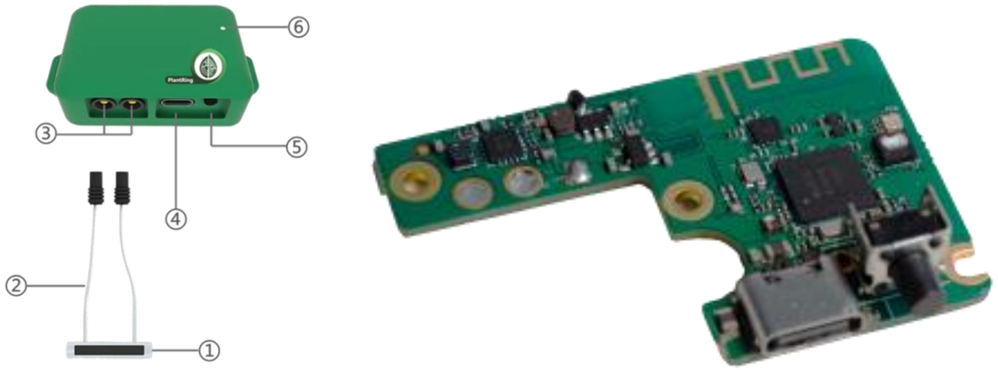


**Fig. S2. Overview of the data logger.** The sensor unit comprises the following components: ① a strain sensor, ② wires with banana plugs, ③ banana sockets, ④ a Type-C interface, ⑤ a power/zeroing button, and ⑥ an LED indicator light. The data logger is rated IPX5 for water resistance, protecting against water projected from a spray nozzle at any angle for 10 to 15 minutes from a distance of 3 meters at a pressure of 30 kPa. The PCB integrates the NRF52833 Bluetooth chip and an ADC module, powered by a dedicated charging management system connected to a 380 mAh battery. It includes operational amplifier circuitry for connecting strain sensors and an air temperature and humidity sensor via an Inter-Integrated Circuit (I2C) protocol. User interaction is facilitated through General-Purpose Input/Output (GPIO) connections, which support an LED light and a multifunction button, enabling operation control, resetting, or shutting down the device. The PCB design also provides for easy battery charging and serial data download via a Type-C interface. Additionally, the PCB is equipped with a PCB antenna, and during operation, the data logger communicates with the gateway using 2.4G RF technology.

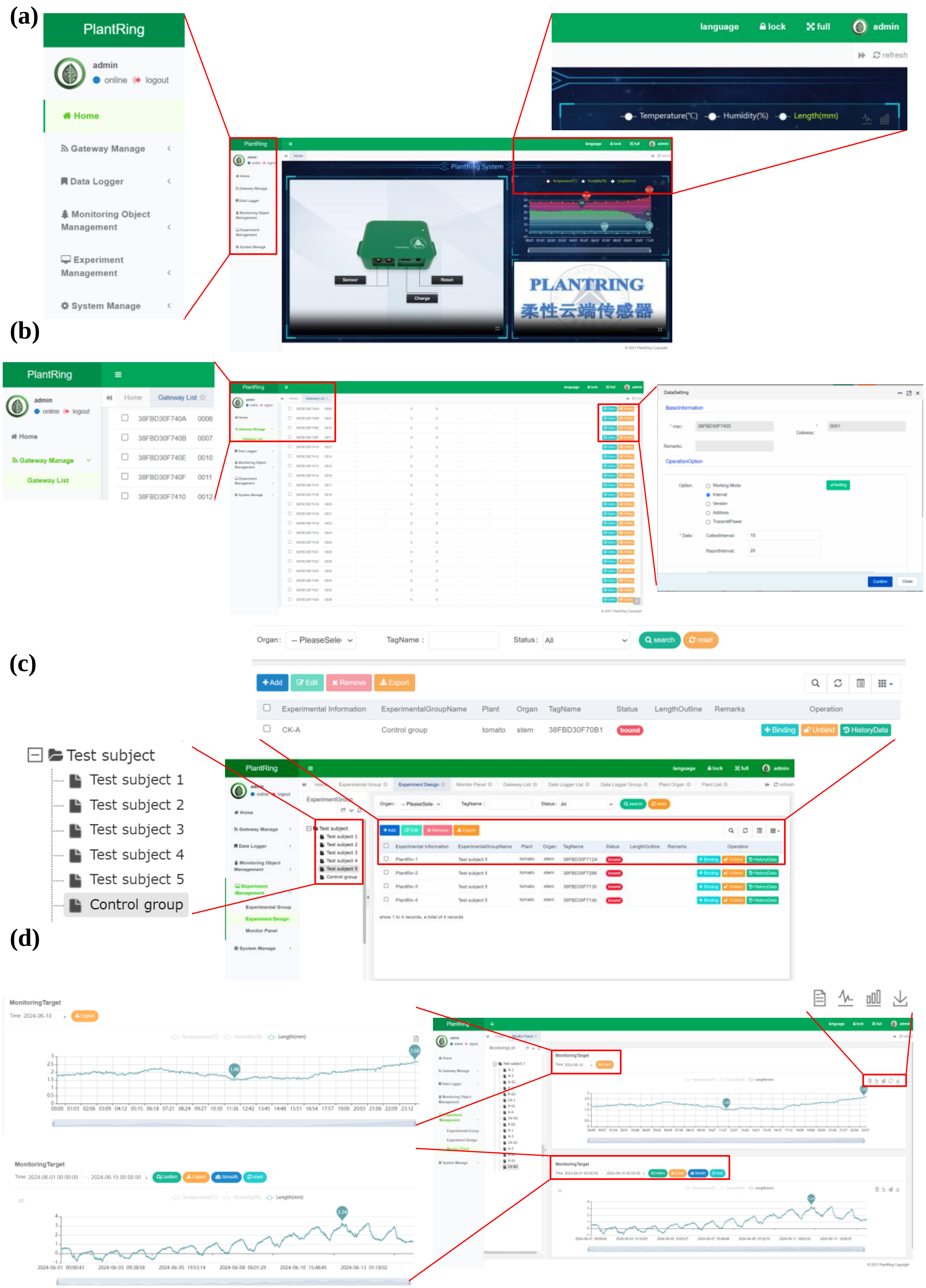


**Fig. S3. Overview of the cloud server interface.** (a) Homepage Layout: The homepage showcases functionalities such as language selection (Chinese and English), full-screen mode, and webpage locking. An example displays shows temperature, humidity, and length values. (b) Gateway Status and Settings: The network currently encompasses over forty gateways, each capable of managing at least 300 PlantRing units. The reporting and sampling intervals for these gateways can be adjusted, as highlighted on the right side of the figure. (c) Sensor Information in Experimental Design: Groups can be created based on experimental parameters to review data from different gateways. The example illustrates how to search and access specific sensor information, including plant species, detection target, sensor ID, and historical data. In this instance, the experimental group is labeled "Control group" with tomato as the plant species, stem as the detection target, and "38FBD30F70B1" as the sensor ID. (d) Data Export for Specific Date Ranges: Users can select a specific date within the timeline to view the 24-hour length curve for that day. By clicking the "Export" button, an Excel file containing time, length values, temperature, humidity data, sensor ID, and battery level can be downloaded. The four icons arranged from left to right offer functionalities for data visualization, line graph representation, bar chart depiction, and curve image preservation. Users can select a start and end date within the timeline, and clicking "Confirm" will display the curve for the chosen date range. The "Smooth" button applies a 20-point smoothing algorithm to the curve, and the "Reset" button allows users to select a new date range.

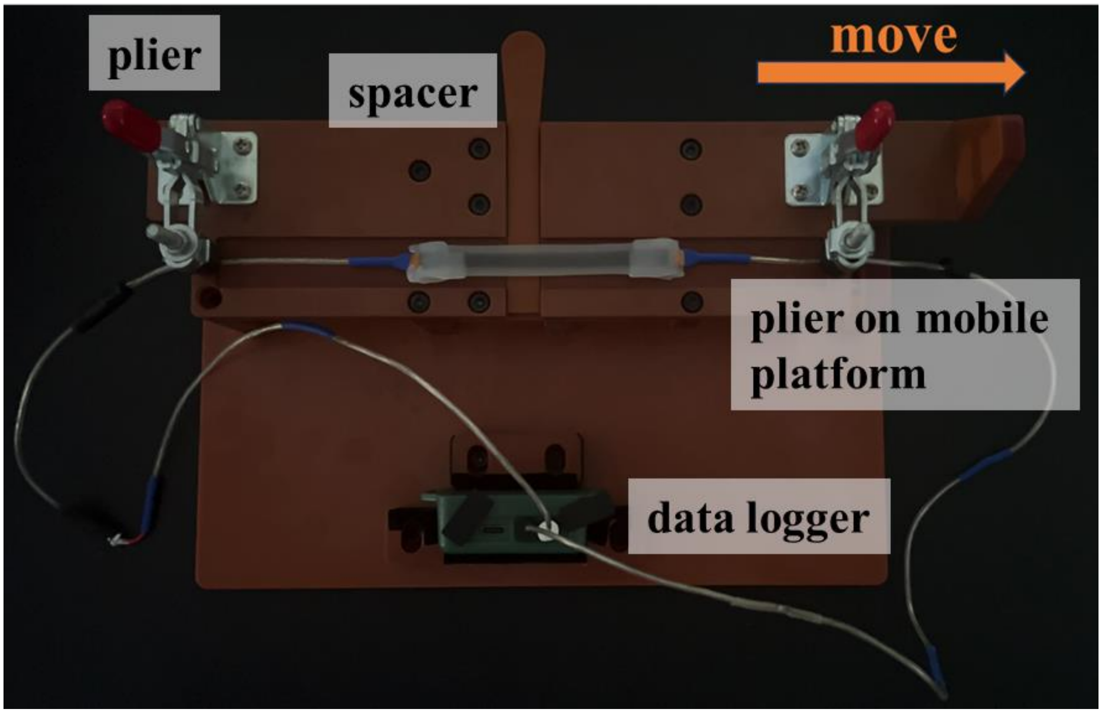


**Fig. S4. The uniaxial measurement device used for signal calibration of sensors.** The device consists of two pliers, one installed on a fixed platform and the other on a mobile platform. The sensor was secured at both ends by the pliers, maintaining its original length at the starting point. The mobile platform can be moved with precise distance along the slide rail to control the strain applied to the sensor. A spacer could be inserted to keep the distance constant. During the stretching process, the sensor was connected to the data logger and the variation of resistance was recorded on the cloud sever. The resistance value was then plotted against the stretched length of the sensor, with the slope of the plot used for signal calibration.

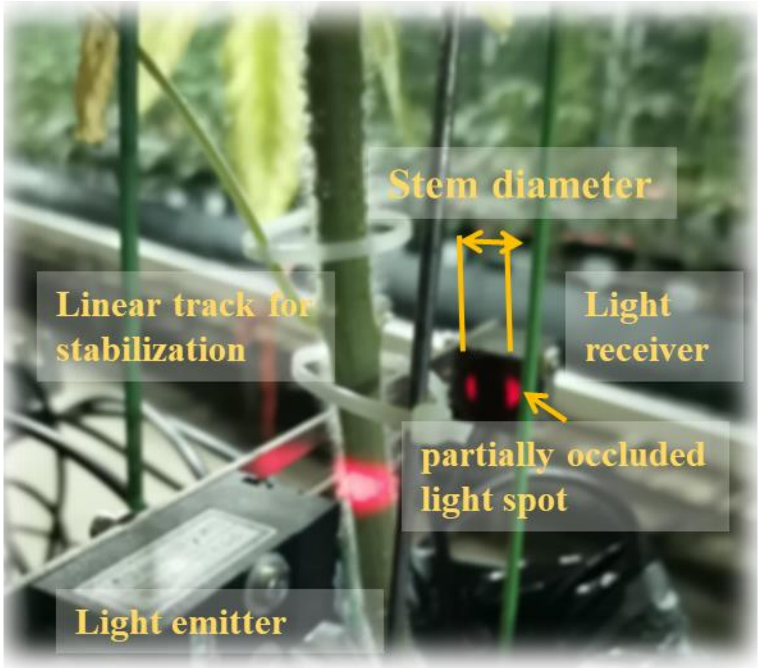


**Fig. S5. Measurement of stem diameter using a laser displacement sensor.** A laser displacement sensor was positioned near the probes at both ends of the PlantRing that was attached to the stem. These sensors were aligned using a custom-designed linear track, ensuring that the stems of the tomato plants consistently remained within the laser beam's range. The light spot was partially occluded by the stem, which could be used to measure the stem diameter with the receiver end.

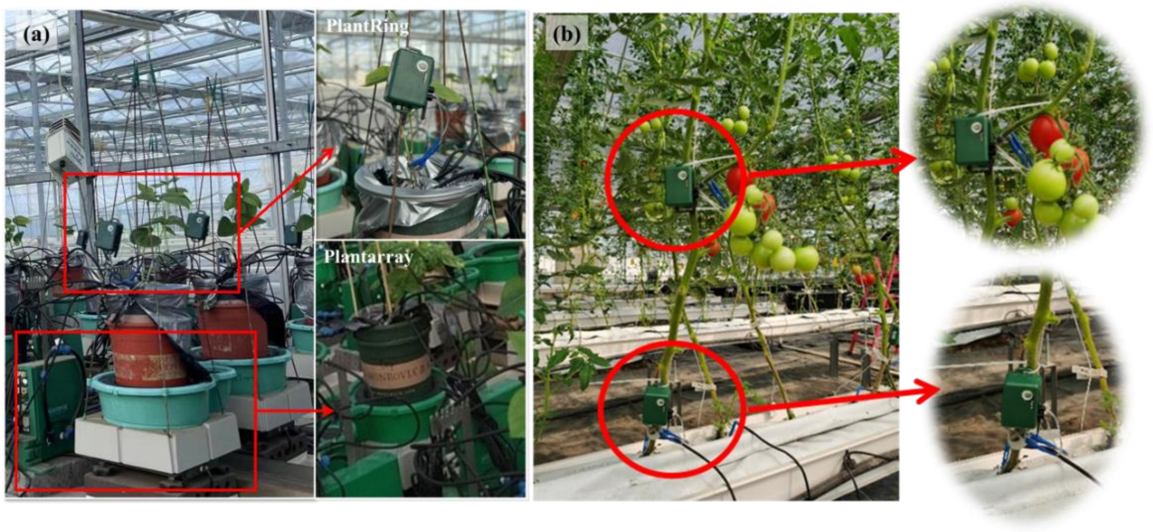


**Fig. S6. Images of plants grown Plantarray and/or equipped with PlantRing.** (a) Common bean plants were cultivated on the Plantarray lysimeter platform and installed with the PlantRing system to compare measurement of water relations. The Plantarray platform dynamically measures the total weight of system, whereas PlantRing system dynamically measures ΔSC of the main stem. (b) Tomato plants each installed with two PlantRing sensors for studying fruit cracking based on the measurement of stem circumferences. One sensor was installed on the main stem, and the other on the fruiting branch.

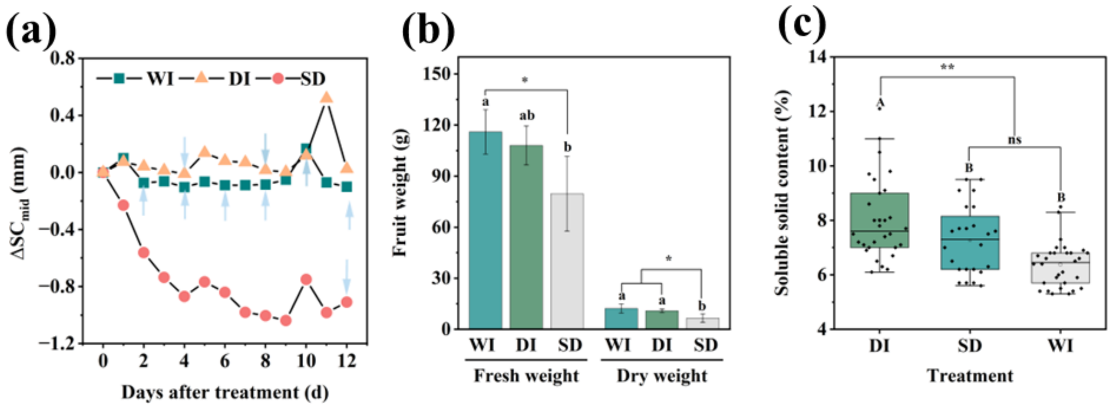


**Fig. S7. Results of PlantRing-based feedback irrigation under laboratory artificial lighting conditions.** The results demonstrate the promising utility of PlantRing for guiding plant-based deficit irrigation as a next-generation approach to achieve water conservation and quality improvement simultaneously. (a) Three irrigation modes were implemented for cultivation of tomato plants, the ΔSC was recorded for comparison. The three modes included: (1) regular irrigation every 2 days (well irrigation treatment, WI), (2) irrigation initiated upon observation of leaf wilting (severe drought treatment, SD), and (3) irrigation initiated when a negative slope was detected using linear regression for the $\Delta SC_{mid}$ over five consecutive days, indicating a significant decrease in $\Delta SC_{mid}$ (deficit irrigation treatment, DI). (b) Fruit productivity under different treatment, measured by the fresh or dry weight of all fruits per plant. The results showed no significant difference between the WI and DI groups, whereas a reduction in yield was noted under SD treatment. A reduced number of fruits per plant (by 26.7%, $P<0.01$), rather than reduced fresh or dry weight of individual fruits ($P>0.05$), was the reason for yield reduction in SD treatment. (c) The measured soluble solid content of fruits, which showed a notable increase in DI compared with WI and SD.

**Table S1. The heating procedure for producing carbonized silk georgette.** The temperature was adjusted from the starting temperature to the heating temperature at different rate, then the temperature was kept constant for different heating time before the next stage starts.

| stage | Starting temperature (℃) | Heating temperature(℃) | Heating/cooling rate(℃/min) | Heating time (min) |
|---|---|---|---|---|
| 1 | 25 | 150 | 10 | 60 |
| 2 | 150 | 350 | 5 | 180 |
| 3 | 350 | 950 | 3 | 120 |
| 4 | 950 | 25 | Natural cool down | / |

**Table S2. Characterization of the repeatability and accuracy of PlantRing system.** The data was obtained from testing 9 different 6 cm type sensor, AVD AD stands for the average AD value measured under different levels of strain, STD stands for the standard derivation of the AD values, CV stands for the coefficient of variation of the AD values, MS stands for measured strain deducted from AD value, which is also displayed and recorded in cloud sever, RE stands for the relative error calculated by comparing measured strain with the actual strain.

| Strain（mm) | AVG AD | STD | CV（%） | MS（mm) | RE（%） |
|---|---|---|---|---|---|
| 0 | 3518.111 | 6.09999 | 0.173388 | 0 | 0.00 |
| 4 | 3443.333 | 3.858612 | 0.11206 | 4.02 | 0.50 |
| 8 | 3322.333 | 15.28252 | 0.459994 | 8.05 | 0.63 |
| 12 | 3204.222 | 21.49304 | 0.670772 | 11.98 | -0.17 |
| 16 | 3081.333 | 4.921608 | 0.159723 | 16.1 | 0.63 |
| 20 | 2973.333 | 7.272475 | 0.24459 | 19.87 | -0.65 |
| 24 | 2857.111 | 1.728483 | 0.060498 | 23.93 | -0.29 |
| 28 | 2739.667 | 1.825742 | 0.066641 | 28.04 | 0.14 |
| 32 | 2622.111 | 7.370277 | 0.281082 | 32.14 | 0.44 |

**Movie S1 (separate file).** The experimental scenario for testing PlantRing under controlled airflow conditions. The tomato plants were placed in a tunnel with natural light and ambient temperatures (ranging from 25 to 34°C on the testing day, November 28th, 2023). Three tomato plants were selected for the experiment, each equipped with a PlantRing attached at its stem base. An oscillating fan (AD61-1, AIRMATE, China) was positioned one meter away from the tomato plants to generate an airflow velocity of approximately 3.5m/s. The wind interference period lasted from 9:00 to 16:00.

**Movie S2 (separate file).** The experimental scenario for testing PlantRing under rainy day. The tomato plants were placed in an open field in a rainy day (December 4th, 2023). Four tomato plants were selected for the experiment, each equipped with a PlantRing attached at its stem base. The rain interference period lasted from 10:00 to 16:00, during which the precipitation rate ranged from 0.1 to 0.38 mm/h.

**Code S1. The code used for temperature compensation.**

The measured rod circumference remained relatively stable below room temperature, and the impact increased with the temperature, which was managed via temperature compensation within the system. The code for temperature compensation is as follows：

```
import os
import numpy as np
import pandas as pd
from sklearn.model_selection import train_test_split
from sklearn.linear_model import LinearRegression
from sklearn.preprocessing import PolynomialFeatures
from sklearn.metrics import mean_squared_error
from sklearn.impute import SimpleImputer


folder_path = r"File Route*"   *:put the local path of data here
files = os.listdir(folder_path)


train_files = np.random.choice(files, 5, replace=False)
print(train_files)


train_data = []
for file_name in train_files:
    file_path = os.path.join(folder_path, file_name)
    data = pd.read_excel(file_path, usecols=[1, 2])
    train_data.append(data)
```

```
train_data = pd.concat(train_data, ignore_index=True)
X_train = train_data.iloc[:, 0].values.reshape(-1, 1)
y_train = train_data.iloc[:, 1].values

imputer = SimpleImputer(strategy='mean')
y_train = imputer.fit_transform(y_train.reshape(-1, 1)).ravel()

imputer = SimpleImputer(strategy='mean')
X_train = imputer.fit_transform(X_train)

test_files = [file_name for file_name in files if file_name not in train_files]

def correct_length(model, X, y):
    return y - model.predict(X)

models = {
    'linear_model': LinearRegression(),
    'quadratic_model': PolynomialFeatures(degree=2),
}

model_errors = {}
for name, model in models.items():
    if name == 'linear_model':
        model_errors[name] = mean_squared_error(y_train, model.fit(X_train,
y_train).predict(X_train))
    elif name == 'quadratic_model':
        poly = PolynomialFeatures(degree=2)
        X_poly = poly.fit_transform(X_train)
        model_errors[name] = mean_squared_error(y_train, LinearRegression().fit(X_poly,
y_train).predict(X_poly))
```

```
best_model = min(model_errors, key=model_errors.get)

if best_model == 'linear_model':
    selected_model = LinearRegression()
    selected_model.fit(X_train, y_train)
elif best_model == 'quadratic_model':
    poly = PolynomialFeatures(degree=2)
    X_poly_train = poly.fit_transform(X_train)
    selected_model = LinearRegression()
    selected_model.fit(X_poly_train, y_train)

for file_name in test_files:
    file_path = os.path.join(folder_path, file_name)
    data = pd.read_excel(file_path, usecols=[1, 2])
    temperature = data.iloc[:, 0].values.reshape(-1, 1)
    length = data.iloc[:, 1].values

    temperature = imputer.transform(temperature)

    if best_model == 'linear_model':
        corrected_lengths = correct_length(selected_model, temperature, length)
    elif best_model == 'quadratic_model':
        poly = PolynomialFeatures(degree=2)
        temperature_poly = poly.fit_transform(temperature)
        corrected_lengths = correct_length(selected_model, temperature_poly, length)
    data[' corrected_lengths '] = corrected_lengths
    output_file_path = os.path.join(folder_path, "output_" + file_name)
    data.to_excel(output_file_path, index=False)

if best_model == 'linear_model':
    print("Linear model coefficient:", selected_model.coef_)
```

```
elif best_model == 'quadratic_model':
    poly = PolynomialFeatures(degree=2)
    X_poly_train = poly.fit_transform(X_train)
    selected_model = LinearRegression()
    selected_model.fit(X_poly_train, y_train)
    print("Quadratic model coefficient:", selected_model.coef_)
else:
    print("The selected model has no coefficient ")
print("The data processing is complete and saved to the output file ")
```
